# Glucocorticoid signaling regulates expression of the EBI3 subunit of IL-27 in neonatal macrophages: Implications for antenatal corticosteroid therapy

**DOI:** 10.64898/2026.03.24.713718

**Authors:** Jordan K Vance, Lei Wang, Jessica M Povroznik, Jonathan T. Busada, Gangqing Hu, Cory M Robinson

## Abstract

**Background:** Humans and mice display elevated levels of IL-27, an immunosuppressive cytokine shown to increase during neonatal bacterial sepsis and compromise survival. This study explores two hypotheses for regulation of IL-27 expression: 1) decreased DNA methylation in newborns that contributes to increased expression of IL-27 genes; 2) neonatal hormones regulate IL-27 expression through upstream hormone response elements (HREs).

**Methods:** Whole genome methyl-seq analysis of neonatal and adult blood-derived macrophages identified differentially methylated regions (DMRs) at steady-state. Quantitative PCR (qPCR) measured expression of IL-27 genes (*IL27p28* and *EBI3)* in human and murine neonatal macrophages stimulated in vitro with synthetic glucocorticoid or progesterone. Confocal microscopy and chromatin immunoprecipitation (ChIP) of glucocorticoid receptor (GR) assessed translocation into the nucleus and binding to the *EBI3* promoter.

**Results:** The IL-27p28 promoter contained DMRs that were increased in the neonatal cohort. The analysis did not identify DMRs within the EBI3 promoter. Dexamethasone stimulation increased *EBI3* gene expression in human and murine neonatal macrophages. GR localized to the nucleus in response to dexamethasone and was enriched at the *EBI3* upstream regulatory region.

**Conclusion:** These data suggest glucocorticoid (GC) signaling increases EBI3 expression. This has importance in the context of antenatal GC administration that may increase IL-27 levels.

**Impact Statement:**

▪ Elevated expression of IL-27 in early life impairs the host response to invasive bacterial infection in neonates.
▪ Understanding the regulatory mechanisms contributing to increased IL-27 during the neonatal period is necessary to reduce susceptibility to infection in this vulnerable population.
▪ The methylation status of the IL-27 genes in macrophages from neonatal and adult blood donors does not suggest regulation of differential expression with age.
▪ Glucocorticoids are a signal that can induce EBI3 gene expression in a GR-dependent manner.
▪ Glucocorticoid therapy for premature infants may increase IL-27 expression and promote enhanced susceptibility to infection.

## 1. Introduction

Infancy is a vulnerable developmental stage with a disproportionately high mortality rate within the first month of life. Infection is a leading cause of neonatal death^1, 2^. Understanding key aspects of development from infancy to adulthood is essential to improving practices in clinical care, as well as the development of appropriate therapeutics to combat disease and infection in specific populations. Suppressive features are well represented in the neonatal immune profile, in part as carryover from the fetal-maternal environment, but also as a mechanism to protect from host-responsiveness to common antigens for the first time. A decreased expression of toll like receptors (TLRs) and reduced Th1 cytokine production in comparison to the adult immune profile, contribute to a Th2-bias and inadequate protective response to microbial pathogens^3–7^. The limitations in combatting early life infections are further exacerbated in cases of preterm (PT) and low birth weight (LBW) infants^8, 9^.

Progesterone and glucocorticoids, steroid hormones with immunosuppressant activities, rise throughout pregnancy to support the developing fetus^10, 11^. Glucocorticoids (GCs) begin to rise through the first trimester, encouraging organ development, fetal adrenal system development, and aiding in overall steroidogenesis^12, 13^. Following delivery, these hormones remain elevated in the newborn for approximately 24 h and up to 96 h, respectively^14, 15^. In addition to these factors influencing the neonatal immune response, we have reported IL-27 as a key factor that regulates the host response to infection in newborns^16, 17^.

IL-27 is a pleotropic cytokine that is comprised of two subunits, p28 and EBI3, which dimerize to signal through the IL-27 receptor made up of gp130 and IL-27Rα. This leads to JAK/STAT signaling through a phosphorylation cascade that activates STAT1 or STAT3 and can promote IFNγ or IL-10 production, respectively. We have demonstrated that IL-27 expression is elevated at baseline in human cord blood-derived macrophages^16, 18^. These observations are consistent in mice that have increased IL-27 transcripts and cytokine-positive cells in the spleens of neonatal and juvenile mice compared to adults^16^. These levels further increase following infection and mice lacking the ability to respond to IL-27 signaling fare better in our established *Escherichia coli-*induced model of neonatal bacterial sepsis^17^. We have curated a detailed profile of IL-27-producing cells during gram-negative neonatal sepsis and following neonatal BCG vaccination; these studies identified discreet subpopulations of myeloid cells as sources of IL-27 production^19, 20^. The overarching premise of our work is that IL-27 neutralization may represent a viable therapeutic approach in the effort to expand the arsenal for clinicians to combat neonatal infections. An essential part of the realization of this goal is to determine the signals and/or regulatory mechanisms that regulate IL-27 expression at baseline and during infection in the neonatal population.

CpG islands reside within the IL-27p28 locus for methylation and hormone response elements (HREs) are located upstream of the IL-27p28 and EBI3 transcriptional start sites. Collectively, these observations led to the hypothesis that epigenetic and hormonal factors may have regulatory roles in IL-27 expression at different periods of life. CpG islands are a series of repeating cytosine and guanine dinucleotides at comparatively high density to which a methyl group can be added to a cytosine base by DNA methyltransferase^21^. Methylation most frequently prevents chromatin remodeling and silences gene expression^22^. Canonically, hormone signaling begins with diffusion through the cell membrane where the free hormone binds available receptor present within the cytoplasmic space. This in turn results in a conformational change that allows the hormone-receptor complex to homodimerize and translocate to the nucleus, binding the appropriate HREs^13, 23–25^. We hypothesized that increased methylation of IL-27p28 into adulthood restricts IL-27 expression, and hormonal influences in the neonatal period contribute to increased IL-27.

We investigated both methylation and hormone signaling as mechanisms that regulate IL-27 expression. While global DNA methylation levels are decreased in cord blood derived macrophages (CBMs) compared to adult peripheral blood macrophages (PBMs), there is greater methylation at the genes encoding IL-27. The glucocorticoid receptor (GR), but not progesterone (PR), is expressed in cord blood mononuclear cells (CBMCs), monocytes, and fully differentiated CBMs. CBMs increase EBI3 gene expression in response to treatment with dexamethasone, a synthetic glucocorticoid (GC) used clinically for antenatal therapy. CBMs demonstrated both increased GR in the nucleus and enriched GR binding upstream of EBI3 with dexamethasone stimulation. Together, our findings provide new insights into signaling pathways and mechanisms that regulate IL-27 gene expression early in life. Furthermore, they have clinical implications when considering standard administration of corticosteroids during pregnancy for anticipated or at-risk preterm labor. As such, this practice could promote precarious levels of IL-27 that increase susceptibility to infection in an already vulnerable population.

## 2. Materials and Methods

### 2.1. Blood donor criteria and macrophage differentiation

All blood donor units were anonymous, deidentified, and obtained with Institutional Review Board approval (protocol no. 2304755320). Fresh human umbilical cord blood was collected from healthy newborns of gestational age ≥37 weeks and obtained from the Cleveland Cord Blood Center (Cleveland, OH, USA). Whole blood was processed for use within 24 h as described previously^18^. Leukocyte-enriched buffy coats from adult donors were purchased from New York Blood Center (New York, NY, USA). Adult donors were healthy individuals that were a minimum of 16 years old and ≥110 lbs. Both male and female sexes were included for both age cohorts. Leukocyte-enriched buffy coats were subjected to density-gradient centrifugation with Ficoll Paque-Plus (Millipore-Sigma, Burlington MA, USA) according to standard procedures to isolate the cord blood or peripheral blood mononuclear cells (CBMCs/PBMCs). Monocytes were isolated from the mononuclear fractions by Optiprep (Millipore-Sigma) density gradient centrifugation as described previously^26^. Cells were cultured for a minimum of 1 h in serum-free Dulbecco’s Modified Eagle Medium (DMEM; Corning, Corning, NY, USA) supplemented with 2 mM glutamine (Cytiva, Marlborough, MA, USA), 25mM HEPES (Cytiva, Marlborough, MA, USA), and 100 U/mL penicillin/streptomycin (Millipore-Sigma). Adherent cells were then cultured with DMEM supplemented with 10% human serum (Fisher Scientific, Fairlawn, NJ, USA), 20% fetal bovine serum (FBS, Avantor, Radnor Township, PA, USA), 2mM glutamine, 25mM HEPES, and 100 U/mL penicillin/streptomycin for 7-10 days to allow differentiation to macrophages.

### 2.2. Animal care and use

Adult C57BL6/J (WT), Nr3c1*^fl/^*^fl^ homozygous for the loxP-flanked (floxed) *Nr3c1* gene, and LysM-Cre^+/+^ or LysM-Cre^+/-^ mice carrying a *Cre* transgene under the control of the LysM promoter on a C57BL/6J genetic background were purchased from Jackson Laboratory (Bar Harbor, ME USA) and maintained under specific pathogen-free conditions in the vivarium at West Virginia University Health Sciences Center. Specific deletion of glucocorticoid receptor (GR) in cells of myeloid-lineage was generated by crossbreeding Nr3c1*^fl/^*^fl^:LysM-Cre^-/-^ and Nr3c1*^fl/^*^fl^:LysM-Cre^+/-^ mice for a predicted 50% distribution of pups that acquire a *Cre* allele. Neonatal pups were defined as ≤8 days of age^16, 27^. Transgenic mice were genotyped by TransnetYX (Cordova, TN, USA). Pup sex was visually determined as previously described^28^. Male and female mice were included in all studies. All procedures were approved by the West Virginia University Institutional Animal Care and Use Committee (protocol no. 1708008935) and conducted in accordance with the recommendations from the Guide for the Care and Use of Laboratory Animals by the National Research Council (NRC, 2011).

### 2.3. Bone marrow-derived macrophage differentiation and spleen harvest

Femurs were collected from 6-8 day neonatal mice as described previously^29^. Bone marrow progenitor cells were counted and seeded at 5×10^5^ −10^6^ cells per well in a multi-well cell culture plate for 5-7 days in L929-conditioned media as described^29^. Prior to stimulation, the media was replaced with that containing 1% FBS overnight to remove possible exogenous signals that could activate GR. Macrophages were stimulated for the designated time with either vehicle (ethanol) or dexamethasone (100 nM; Steraloids, Newport, RI). RNA was extracted as described below. From the same mice used for bone marrow collection, spleens were collected in 500 µL of TriReagent and stored at - 80°C.

### 2.4. Methylation studies

Neonatal or adult macrophages were treated with or without 5-azacitidine (50 μM) for 24 h prior to RNA extraction. For whole genome methylation analysis, genomic DNA (gDNA) was extracted from a minimum of 10^6^ macrophages using the Monarch Genomic DNA Purification kit (product no. TS3010S, New England Biolabs, Ipswich, MA, USA) per manufacturer recommendations. Enzymatic conversion of unmethylated cytosine bases with the NEBNext Enzymatic Methyl-Seq kit (New England Biolabs, Ipswich, MA, USA) and library preparation was performed by the WVU Genomics Core. Sequencing was performed by Admera (South Plainfield, NJ, USA) using an Illumina NovaSeq S4. In total, 9 neonatal samples and 11 adult samples were used for downstream analysis.

### 2.5. Methyl-seq analysis

Paired-end Enzymatic Methyl-seq (EM-seq) reads from enzymatically-converted genomic DNA from human neonatal or adult macrophages blood donors were aligned to the human reference genome (hg38) using bs_seeker2-align.py (BS-Seeker2 v2.1.8) with the Bowtie2 backend^30^. The hg38 reference genome was bisulfite-converted and indexed using the bs_seeker2-build.py script. Duplicate reads were removed prior to downstream methylation calling. Methylation levels at each cytosine site were subsequently extracted using bs_seeker2-call_methylation.py, generating CGmap and ATCGmap output files for use in methylation analysis with CGmapTools^31^. Differentially methylated regions (DMRs) were identified using **CGmapTools** with a dynamic fragmentation strategy, where adjacent CpG sites (≤100 bp apart) were grouped into fragments (maximum length = 1000 bp, minimum = 5 CpGs), and regions showing significant methylation differences (*p* ≤ 0.05, |ΔmC| ≥ 0.1, *t*-test) between adult and neonatal macrophages were defined as DMRs. Genomic annotation and CpG island information were retrieved from the DBCAT database^32^. Representative DMRs located near the IL-27 and EBI3 loci were visualized in Integrative Genomics Viewer (IGV)^33^.

### 2.6. Hormone stimulations

Macrophages were cultured overnight in 1% serum-containing media supplemented as described above prior to hormone stimulation. Cells were treated with progesterone (0.1-10 μg/mL; Millipore Sigma) or dexamethasone (1-100 nM; Steraloids) for 2-48 h as described in the figure legends. In experiments that included RU486 (1 µM; Steraloids), cells were pre-treated for 1 h prior to stimulation. All hormones and hormone analogs were dissolved in pure molecular biology-grade ethanol and filter sterilized prior to use. Vehicle treatments were performed with filter-sterilized ethanol diluted in the same manner as the corresponding treatment.

### 2.7. RNA extraction and quantitative PCR gene expression analysis

Macrophages were harvested in TriReagent (Millipore-Sigma) and RNA was extracted using the Zymo Direct-Zol kit (product no. R2061, Irvine, CA, USA). Frozen spleens collected in TriReagent as described above were thawed and homogenized prior to RNA extraction using a modified protocol of the Omega EZNA HP Total RNA kit (product no. R6812-01, Norcross, GA, USA). Chloroform (30 µL; VWR, Radnor, PA, USA) was added to the homogenate and incubated at room temperature for 15 min followed by centrifugation at 15,000×g for 15 min at 4°C. The upper aqueous layer was removed, mixed with an equal volume of 70% molecular grade ethanol (Fisher Scientific), and applied to the spin column. Subsequent steps were performed according to manufacturer protocol and the eluted RNA was quantified using a Nanodrop (ThermoFisher Scientific, Waltham, MA, USA). RNA (500 ng −1 µg) was converted to first-strand cDNA using the Bio-Rad iScript cDNA synthesis kit (product no. 1708890, Hercules, CA, USA) according to manufacturer instructions. cDNA was diluted 1:3 with molecular biology grade water. Gene-specific primer probe assays for gene expression studies are outlined in Table 2.1.

**Table 2.1.**
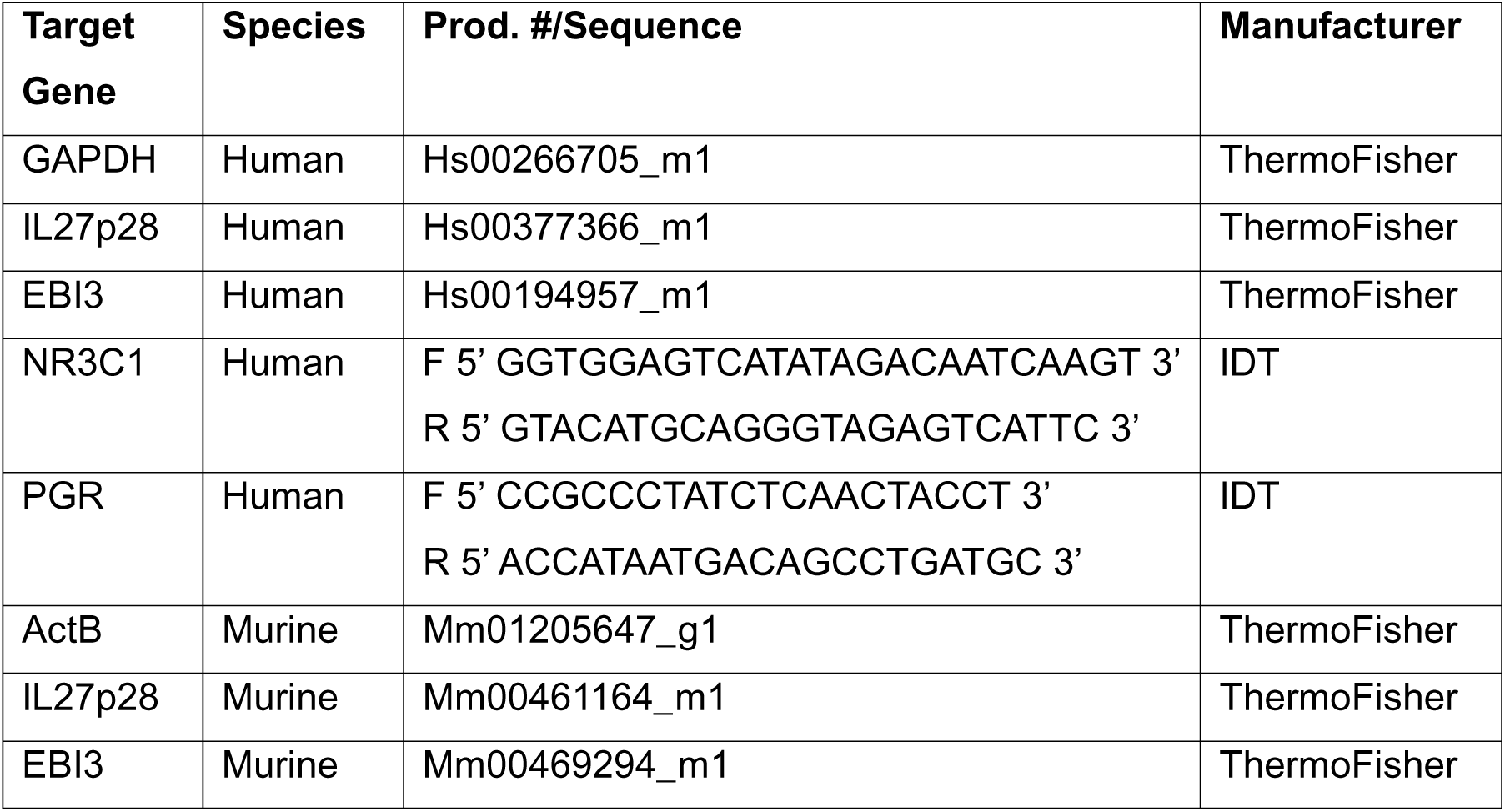
Primer information for gene expression studies.

Real-time cycling reactions that contained diluted cDNA, appropriate primer probes, and iQ Supermix (Bio-Rad) were prepared in triplicate and analyzed using an Applied Biosystems (Waltham, MA, USA) StepOne Plus detection system. Amplification was normalized to internal reference genes glyceraldehyde phosphate dehydrogenase (GAPDH, human) or β-actin (ActB, mouse) and expressed as the log_2_ relative change in gene expression compared to the vehicle controls using the formula 2^-ΔΔCt^. Gene expression levels of PGR and GR were normalized to GAPDH and expressed as ΔCt values. When comparing p28 and EBI3 levels between WT^fl^ and GRKO spleens, all samples were normalized to the lowest expressor within the respective gene.

### 2.8. Immunofluorescent imaging and analysis

Human neonatal macrophages were differentiated as described above on fibronectin-coated coverslips in a 24-well plate and stimulated as described. At the conclusion of the timepoint, cells were fixed in a final concentration of 2% paraformaldehyde (PFA; ThermoFisher Scientific, Waltham, MA, USA) for 15 min at room temperature. Cells were washed and permeabilized in 0.1% Triton X-100 (ThermoFisher Scientific, Waltham, MA, USA) for 10 min at room temperature and subsequently blocked in PBS that contained 1% bovine serum albumin (BSA, Fisher Scientific, Fairlawn, NJ, USA) for 10 min. Rabbit polyclonal GR antibody (1:3200, #3660, Cell Signaling, Danvers, MA, USA) and phalloidin (1:1000; AlexaFluor-594, Life Technologies, Carlsbad, CA, USA) were incubated overnight at 4°C. Cells were washed in PBS that contained 1% BSA and incubated for 30 min with anti-rabbit IgG AlexaFluor-488 (1:4000, Life Technologies). After additional wash steps, coverslips were mounted with Prolong Diamond with DAPI antifade mountant (Invitrogen, Waltham, MA, USA) and cured at room temperature overnight. Confocal microscopy imaging was performed using a Zeiss 710 (Oberkochen, Germany) at 63X magnification. Images within an experiment were analyzed using Fiji (v. 2.16.0)^34^. Analysis of GR translocation to the nucleus was performed with CellProfiler (v. 4.2.8)^35^. The analysis pipeline first split the channels (ColorToGray), identified nuclei from the DAPI channel (IdentifyPrimaryObjects), and merged nuclei within 5 pixels (SplitOrMergeObjects). Signal measurements from the red (actin) and green (GR) channels were merged to calculate the maximum pixel intensity within the cell boundary (ImageMath). Using the previously defined nuclei and cell boundary, the whole cell area was calculated (IdentifySecondaryObjects) and the cytoplasmic signal intensity was subtracted from the whole cell (IdentifyTertiaryObjects). The difference of intensity was calculated to quantify colocalized nuclear and GR signal (MeasureObjectIntensity). For each individual cell, the nuclear mean intensity was divided by the mean intensity of the cytoplasm (CalculateMath).

### 2.9. Chromatin immunoprecipitation

Human neonatal macrophages were seeded at 1-5×10^6^ cells per well in a 24- or 6-well plate. Cells were stimulated with vehicle or dexamethasone (100 nM) for 24 h. Chromatin isolation and immunoprecipitation was carried out as previously described^18^.

Cells were crosslinked in 2% formaldehyde for 2 min at room temperature, quenched with glycine (0.1 M), and the cells were washed in PBS. Cells were scraped on ice, collected in the residual PBS, and flash frozen on dry ice. The solution was thawed with 100 μL of Buffer C that contained 20 mM HEPES pH 7.9, 25% glycerol, 0.42 M NaCl, 1.5 mM MgCl_2_, and 0.2 mM EDTA. Nuclei were pelleted at 10,000×g for 10 min at 4°C. Nuclei were resuspended in 25 µL of breaking buffer that contained 50 mM Tris-HCl pH 8.0, 1 mM EDTA, 0.15 M NaCl, 1% SDS, and 2% Triton-X 100. Chromatin was sonicated twice for 20 s separated with a 30 s rest on ice. The fragmented chromatin was electrophoresed on 1.5% agarose to normalize the amount of nucleic acid by band intensity analysis using a ThermoFisher Scientific iBright 1500 (Waltham, MA, USA). Equal amounts of chromatin were used for immunoprecipitation with the Abcam One Step ChIP Kit (product no. ab117138, Cambridge, UK). Monoclonal anti-GR (0.4 µg/mL, product no. 3660, Cell Signaling, Danvers, MA, USA) was used to pull-down chromatin bound to glucocorticoid receptor. Primers were designed to amplify a 137 bp sequence from publicly available data in the UCSC Genome browser^36^ that showed binding of GR in the upstream regulatory region of *ebi3*^37^ (Supplemental **Error! Reference source not found.**B) and synthesized by Integrated DNA Technologies (IDT, Coralville, IA, USA). The EBI3 promoter region primer sequences were 5’-TTCTCTGTCTCTCTGCTC-3’ (forward) and 5’-TTCCCAGCACAGCATGTC-3’ (reverse). Visualization of the location of the primer (hg38_dna range=chr19:4229086-4229387) is shown in Supplemental **Error! Reference source not found.**B. Real-time cycling reactions that contained equal amounts of eluted DNA, SsoFast EvaGreen Supermix (product no. 1725201, Bio-rad), and 500 nM of primer in triplicate was performed using the Applied Biosystems StepOne Plus detection system. Cycle threshold values were background corrected to no template controls and expressed relative to the non-immune serum immunoprecipitated samples using the formula 2^-ΔΔCt^.

### 2.10. Statistics

Statistical significance was determined with the appropriate parametric or non-parametric test for each of the datasets as described in the figure legends. Data were analyzed as mean ± SEM using unpaired t-tests, Mann Whitney tests, and/or two-way ANOVAs. Statistical significance was analyzed using GraphPad Prism 10 (Boston, MA, USA). The threshold for significance was set to alpha=0.05.

## 3. Results

### 3.1. Neonatal macrophages display lower levels of global methylation compared to adult macrophages

It is well established that patterns of epigenetic modifications change with age and development due to heritable factors, environment, and maturatity^38, 39^. Given the presence of CpG islands in and near the IL-27p28 locus, we hypothesized that differences in IL-27 expression from infancy to adulthood, may be regulated by methylation levels. This hypothesis was tested using a whole genome approach to compare methylation patterns between macrophages differentiated from neonatal and adult blood monocyte donors. The methylation status was determined by comparing changes between the reference genome and enzymatically converted samples at CpG islands in the adult cohort to that in neonates. This identified 2,018 sites with increased and 800 sites with decreased methylation across the genome in adult relative to neonatal donors (Figure 5.1A). These results supported the expectation of elevated global methylation in adult donor macrophages. As such, we evaluated whether or not inhibiting methylation in adult donors would promote increases in expression of IL-27 genes. Neonatal and adult macrophage donors were treated with the DNA methyltransferase (DMT) inhibitor, 5-azacitidine. Neonatal macrophages did not increase expression of IL-27p28 relative to untreated controls (Figure 5.1B). In contrast, adult macrophage donors increased IL-27p28 gene expression to levels significantly greater than those observed in neonates (Figure 5.1B). Treatment with 5-azacitidine did not significantly alter gene expression of EBI3 in either cohort (not shown). Collectively, these data suggested that methylation of sequences within or around the IL-27p28 locus may limit IL-27p28 expression in adult cells.

**Figure 5.1.**
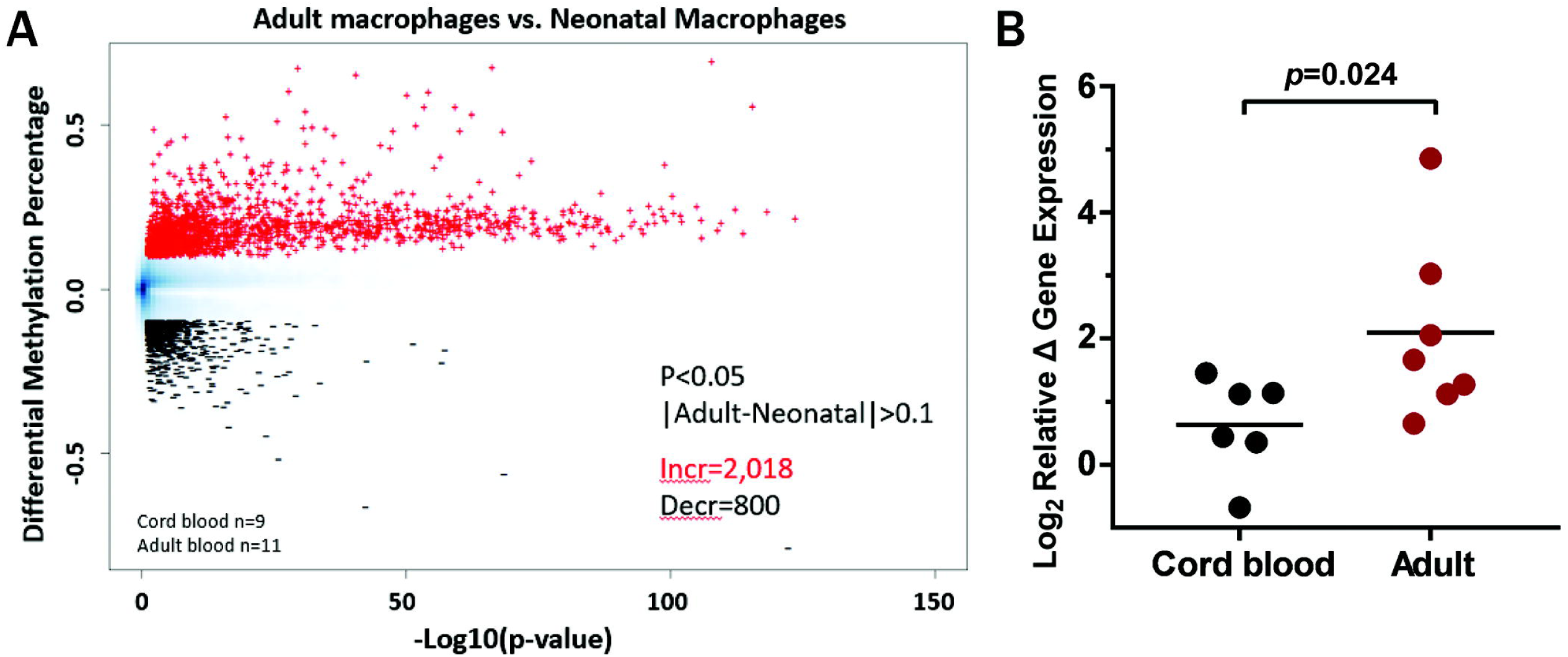
Overall methylation is increased in adult macrophages. (**A**) Macrophages differentiated from cord blood (n=9; black) or adult peripheral blood (n=11; red) were harvested for gDNA that was enzymatically converted to compare differences in methylation status across the entire genome. Percentage of differential methylation with the neonatal population serving as the baseline was found increased in the adult population in 2,018 regions and decreased in 800 regions (p<.05, |Adult-Neonatal| >0.1). (**B**) Macrophages from cord (n=6; black) or adult peripheral blood (n=7; red) were treated with 5-azacitidine (50 μM) for 24 h. Mean log_2_ gene expression levels of IL-27p28 ± standard error of the mean (SEM) determined relative to untreated control cells by real-time PCR using the formula 2^-ΔΔCt^ are shown.

### 3.2. Methylation at the regulatory region of the IL-27p28 gene is reduced in adult macrophages

To explore a methylation-specific regulation of IL-27, a detailed analysis of the promoter regions of IL-27p28 and EBI3 genes was performed. When comparing differentially methylated regions (DMRs) that corresponded to a CpG island within the promoter region of IL-27p28 between neonatal and adult macrophages, we observed a statistically greater degree of methylation in neonatal macrophages (Figure 5.2A-B).

**Figure 5.2.**
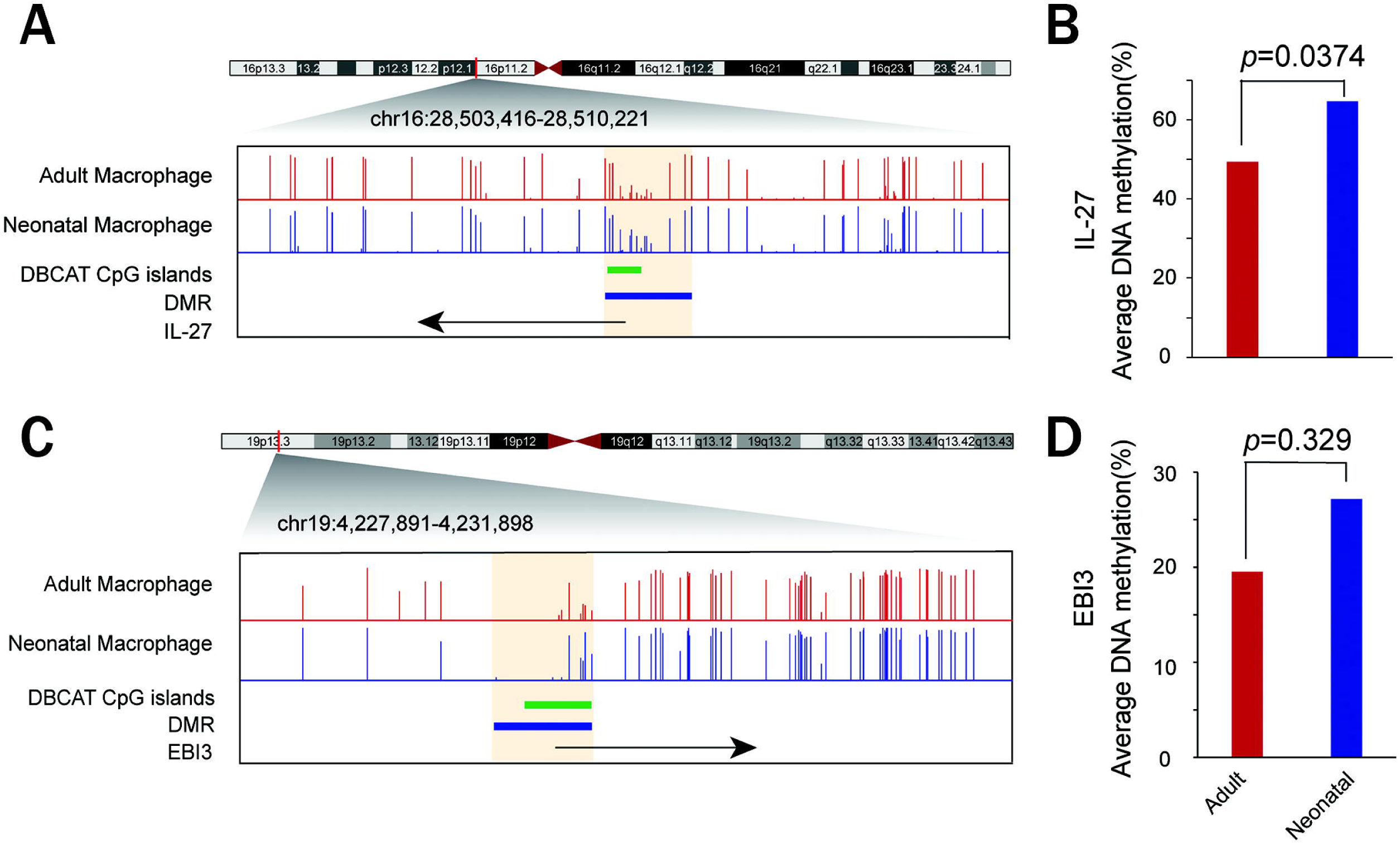
Methylation at the promoter region of the IL-27p28 gene is reduced in adult macrophages. Normalized sequencing read density for adult (red) and neonatal (blue) macrophages at the (**A**) IL-27p28 and (**C**) EBI3 loci indicated are shown. Methylation levels were calculated as the average CpG methylation across all sites at CpG islands (green) within the differentially methylated regions (DMRs) of the promoter for (**B**) IL-27p28 and (**D**) EBI3. Statistical significance was determined using unpaired t tests; exact P values are shown.

Analysis of EBI3 did not show significantly different changes in methylation between neonatal and adult macrophages, suggesting that methylation does not specifically influence EBI3 expression (Figure 5.2C-D). Gene expression analysis of the represented donors showed that IL-27p28 was elevated in the neonatal population, consistent with prior results (data not shown)^16^. Overall, these data suggest that the decrease in IL-27 expression from infancy to adulthood is not the direct result of methylation-related changes to the genome and suggest a separate regulatory mechanism.

### 3.3. Neonatal macrophages express glucocorticoid receptor that translocates to the nucleus following stimulation with dexamethasone

Hormones like progesterone and GCs provide essential signaling for healthy fetal development and progression of pregnancy. Pregnancy is sustained by increasing progesterone until delivery when a sharp drop signals the start of labor^14, 40, 41^. GCs begin to rise at the start of pregnancy and remain elevated to aid in organ development and maturation in the fetus; an additional sharp increase occurring in the final days of gestation prepares the lungs for the extrauterine environment^13, 15, 42, 43^. In silico analysis of the IL-27p28 and EBI3 regulatory regions identified both progesterone response elements (PREs) and glucocorticoid response elements (GREs), respectively. In our prior work, IL-27 gene expression was increased in adult human macrophages exposed to high levels of progesterone, although the change was only significant for EBI3^16^. Additional publications have addressed the influence of GCs on regulation of the hypothalamus-pituitary-adrenal (HPA) axis and steroidogenesis in utero and during the neonatal period^13^. To begin to evaluate the possibility that either progesterone or GC signaling influences neonatal levels of IL-27, we first measured expression of both progesterone receptor (*PGR*) and GR (*NR3C1*) genes in different cellular sources that include or contribute to macrophage development. We found that PGR expression was not detectable in cord blood mononuclear cells (CBMC), monocytes, or monocyte-derived macrophages (Figure 5.3A). In contrast, the GR gene was expressed in all of these cell populations (Figure 5.3B).

**Figure 5.3.**
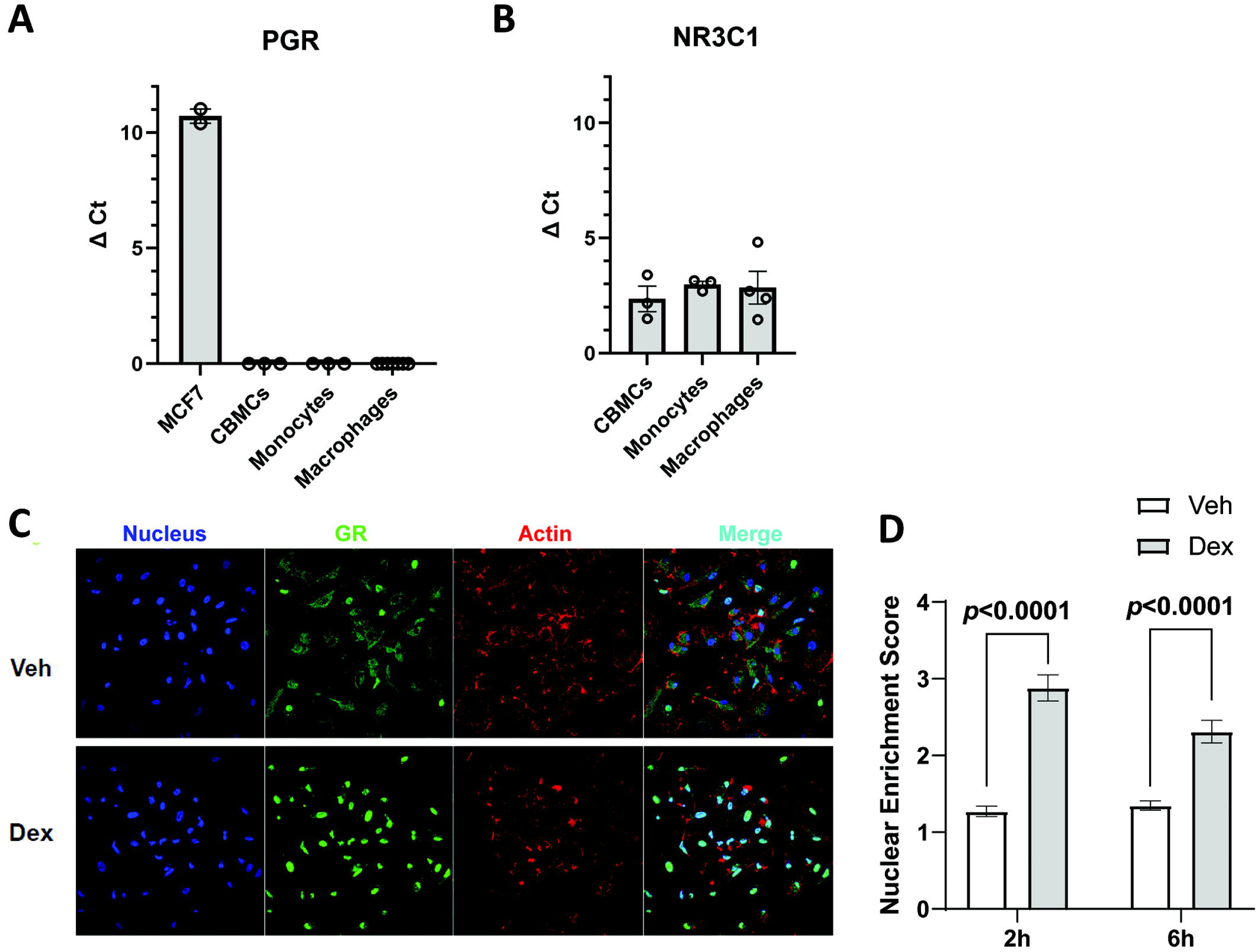
Neonatal macrophages express the glucocorticoid receptor that localizes to the nucleus in neonatal macrophages following exposure to dexamethasone. (**A-B**) RNA was isolated from the cell type indicated at steady-state. The expression of (**A**) progesterone receptor (PGR) and (**B**) glucocorticoid receptor (GR, Nr3c1) was determined by real-time PCR and represented as the change in cycle threshold (Δ Ct) relative to the GAPDH internal control. MCF7 breast cancer cells were used as a positive control for PGR expression. Cord blood mononuclear cells (CBMCs; n=3), monocytes (n=3), and cord blood-derived macrophages (n=5). (**C**) Neonatal macrophages were stimulated with dexamethasone (100 nM) for 2 (n=3) or 6 h (n=5) and immunofluorescent microscopy was performed as described in *Methods* to visualize the nucleus (blue), GR (green) and F-actin (red). Cells were cultured in charcoal-stripped 1% serum-containing media overnight prior to stimulation to limit possible off-target background signaling. Representative images are shown. (**D**) The ratio of GR present in the nucleus was calculated as described in *Methods*. Statistical significance in the 95% confidence interval was determined using one-way ANOVA.

To further confirm GR expression at the protein level, as well as localization to the nucleus in response to GCs in neonatal macrophages, we performed immunofluorescent microscopy. Neonatal macrophages were stimulated with dexamethasone for 6 h and labeled for GR (green) relative to the F-actin cytoskeleton (red) and nucleus (blue, Figure 5.3C). The fluorescent signal from imaging experiments was quantified to compare enrichment of GR in the nucleus following stimulation. This analysis was performed at 2 and 6 h post-stimulation to ensure optimal visualization of GR in the nucleus. Colocalization of GR in the nucleus was evident at baseline with a significant increase in overlapping signal in the dexamethasone-treated cells compared to vehicle controls at each time point (Figure 5.3D).

### 3.4. Glucocorticoids induce expression of EBI3 in neonatal macrophages

Since GR expression was confirmed in all CBMCs, we treated neonatal macrophages with increasing concentrations of dexamethasone or progesterone for 48 h and measured IL-27p28 and EBI3 gene expression (Figure 5.4A-B). Consistent and statistically significant increases in gene expression from the vehicle control were only observed for EBI3, and in response to dexamethasone at 100 nM (Figure 5.4B). RU486, also known as mifepristone, competitively binds PR and GR over progesterone or GCs and antagonizes both PR and GR. This compound is used clinically in the termination of early pregnancy as it prevents the surge in both progesterone and GC necessary to maintain and grow a fetus^44^. In vitro, pretreatment with RU486 prior to neonatal macrophage stimulation with dexamethasone prevented increased EBI3 gene expression (Figure 5.4C-D). Collectively, these findings support the hypothesis that GC signaling through GR regulates EBI3 gene expression.

**Figure 5.4.**
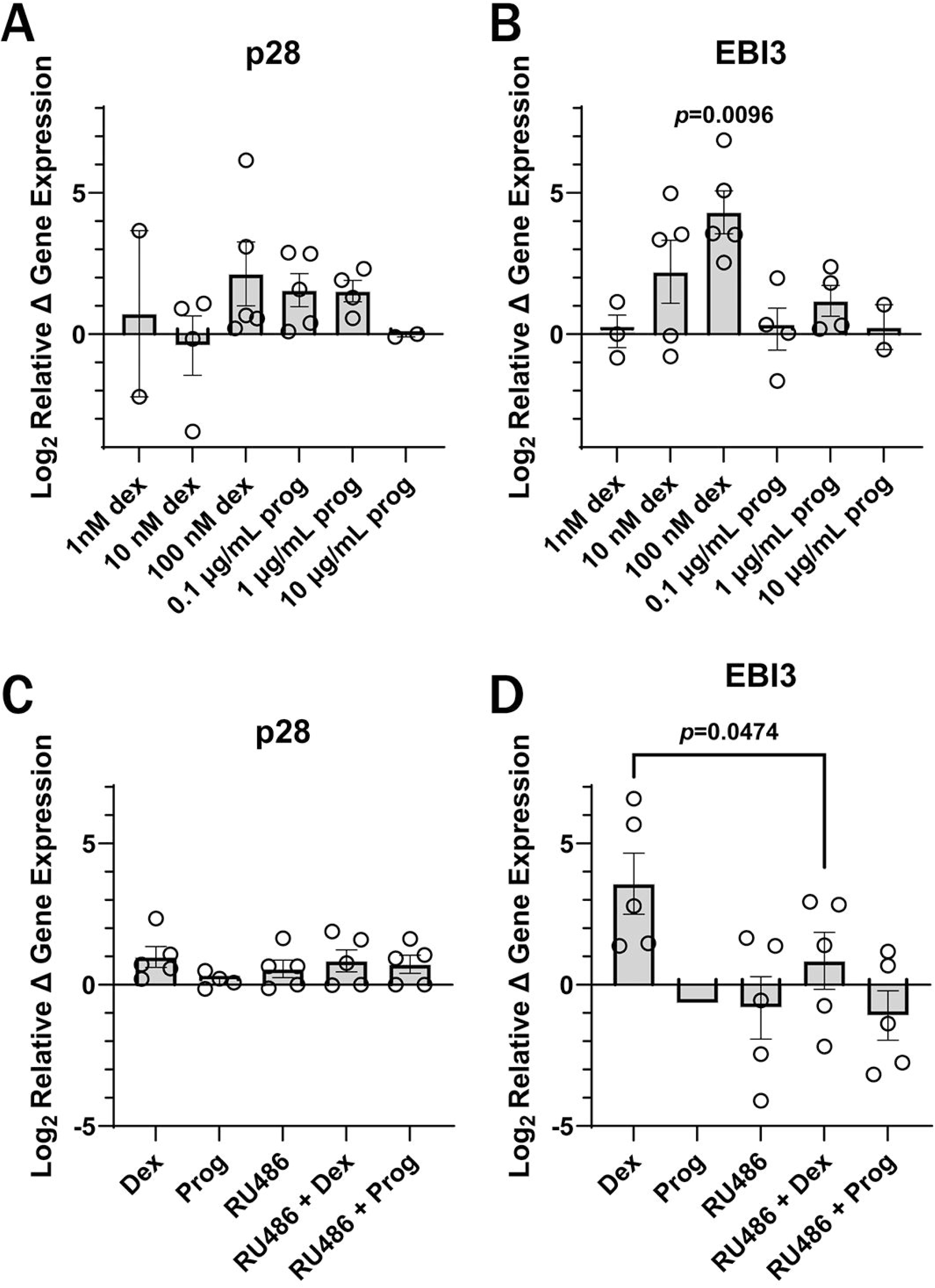
Neonatal macrophages respond to dexamethasone in a dose-dependent and GR-dependent manner to increase EBI3 gene expression. (**A-B**) Neonatal macrophages were treated with the concentration of dexamethasone or progesterone indicated for 48 h. Cells were cultured in 1% serum-containing media overnight prior to stimulation to limit possible off-target background signaling. Mean gene expression levels ± SEM for (**A**) p28 and (**B**) EBI3 for 2-5 independent blood donors are shown. Each symbol represents data from a separate blood donor. (**C-D**) Neonatal macrophages were treated for 1 h with vehicle or RU486 (1 μM) prior to stimulation with vehicle, progesterone (0.1 μg/mL), or dexamethasone (100 nM) for 48 h. Cells were cultured in 1% serum-containing media overnight prior to stimulation. Mean gene expression levels ± SE for (**C**) p28 and (**D**) EBI3 were determined by real-time PCR from 5 independent donors. Gene expression values were normalized to the GAPDH endogenous control and expressed as the log_2_ change relative to vehicle controls using 2^−ΔΔCt^. Statistical significance in the 95% confidence interval was determined by a one-way ANOVA.

### 3.5. GR binds within the promoter region of EBI3 in response to dexamethasone

Since GR translocated to the nucleus (Figure 5.3) and induced expression of EBI3 (Figure 5.4), we sought to determine if this was a product of direct gene regulation. To measure direct recruitment of GR to the EBI3 regulatory region, we performed ChIP analysis. In silico analysis of the EBI3 promoter region identified a consensus GRE sequence that was shown as bound to GR in publicly available ChIP-Seq results^37^. Visualization of publicly available data and primer design for the amplicon of interest are available in Supplemental **Error! Reference source not found.**. Neonatal macrophages were treated with vehicle or dexamethasone for 24 h and chromatin was harvested for ChIP-qPCR analysis of the reported sequence. Comparison of vehicle and dexamethasone treatment groups showed increased enrichment of GR in the dexamethasone group (Figure 5.5).

**Figure 5.5.**
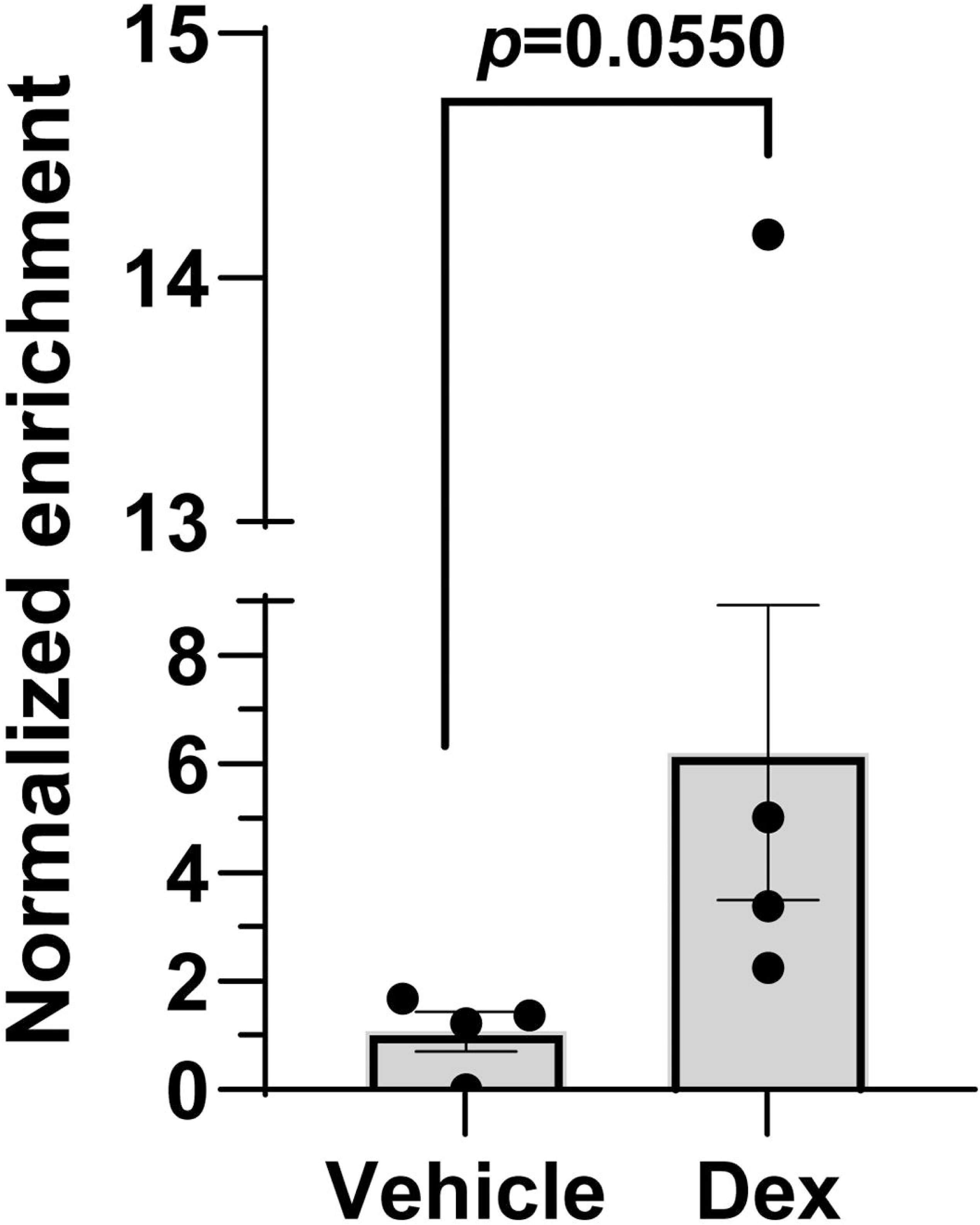
Chromatin immunoprecipitation demonstrated enrichment of GR in the regulatory region of EBI3. Neonatal macrophages were stimulated for 24 h with either vehicle or dexamethasone (100 nM). Cells were cultured in 1% serum-containing media overnight prior to stimulation to limit possible off-target background signaling. Chromatin was isolated as described in the *Methods* and sheared by sonication. GR-bound DNA was immunoprecipitated and amplification of a 137 bp segment that contains a GRE upstream of the EBI3 transcriptional start site was performed by real-time PCR. The Ct values were normalized to no template controls and expressed relative to non-immune serum immunoprecipitated controls according to the formula 2^-ΔΔCt^; shown as normalized enrichment are data from 4 independent donors. Statistical significance was determined in the 95% confidence interval using a two-sample t-test.

### 3.6. Elevated levels of IL-27 in early life are not exclusively regulated by glucocorticoid signaling

We established GC signaling as a mechanism that drives EBI3 gene expression, so we sought to determine if this regulatory axis was responsible for levels of IL-27 that are elevated at baseline in early life. Treatment of murine neonatal BMDMs with dexamethasone yielded results similar to those observed in human macrophages. This allowed us to utilize genetic tools for modeling additional findings in mice (Supplemental **Error! Reference source not found.**). GR(Nr3c1)^fl/fl^ mice were bred to LysMcre^+/+^ to generate GR^fl/fl^LysMcre^+/-^ (GRKO) pups. These mice do not express GR within the myeloid population. We hypothesized that EBI3 expression levels in GRKO mice would be lower in comparison to LysMcre^-/-^ (WT) littermates at baseline due to the lack of GR signaling. As such, we measured gene expression in the spleens of 7-8 days old neonatal mice. The splenic gene expression results did not show a significant difference in IL-27p28 or EBI3 gene expression across genotypes (Figure 5.6A-B). We considered the possibility that detectable differences in myeloid cells may be difficult to visualize in the mixed cell population of the spleen and generated an enriched population of neonatal macrophages derived from bone marrow progenitors of WT and GRKO mice. At baseline levels, we did not observe a difference in expression of either IL-27 gene within the macrophage population (Figure 5.6C-D). This finding suggested that although GC signaling can upregulate EBI3 expression, steady-state IL-27 levels in neonates were not controlled by endogenous GCs.

**Figure 5.6.**
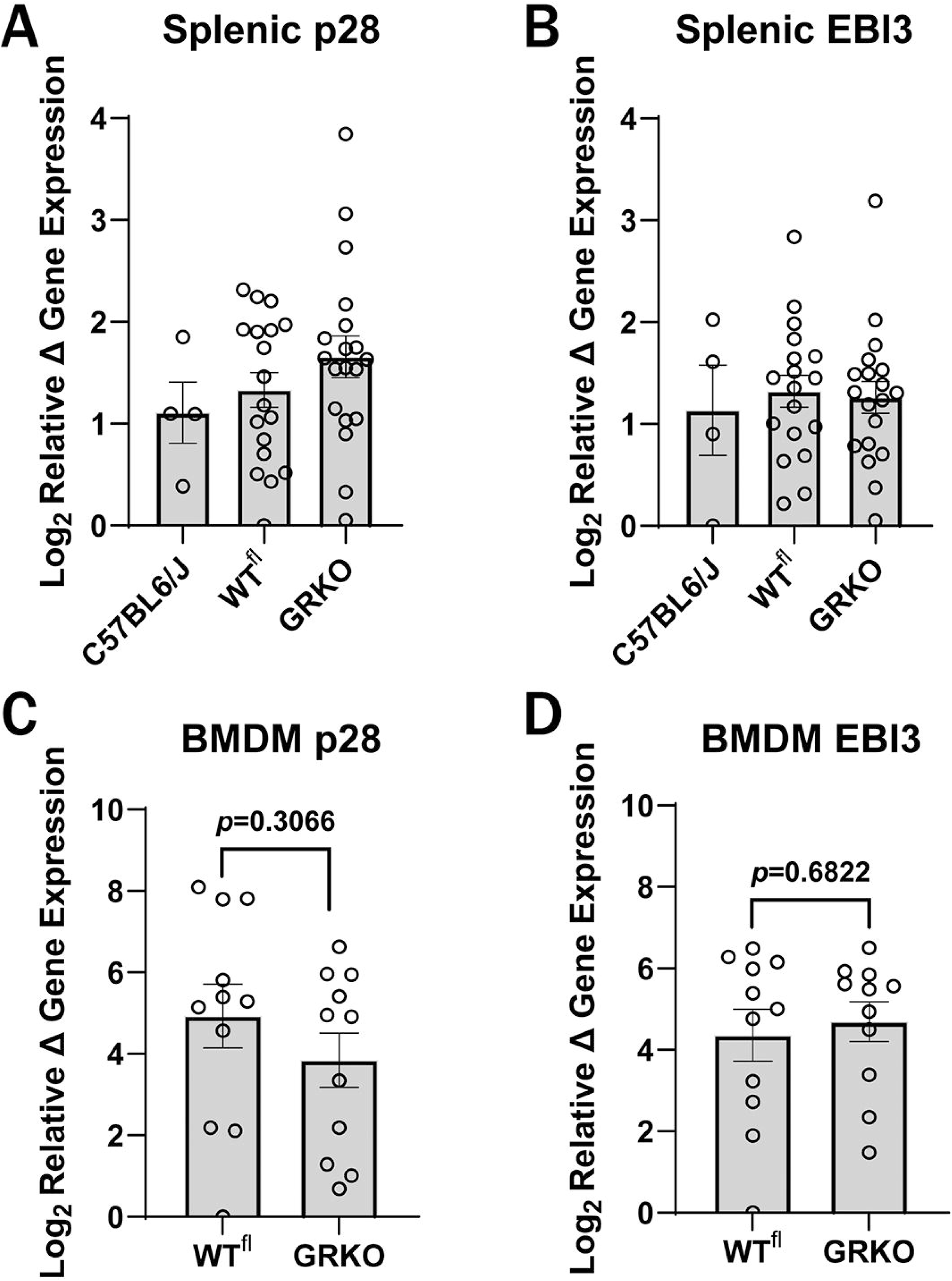
IL-27 gene expression of the IL-27 genes from wildtype or myeloid-specific GRKO mice did not show significant changes in the spleen or BMDMCs. (**A**-**B**) IL-27p28 and EBI3 gene expression profiles from the spleens of neonatal C57BL6/J (n=4), WT^fl^ (n=17), and myeloid-specific GRKO (n=17) mice harvested at baseline on day 7 of life. (**C**-**D**) Bone marrow was harvested from neonatal pups on day 7 of life and macrophages were differentiated from bone marrow progenitors as described in the *Methods*. BMDMs (n=11) were cultured in 1% serum-containing media overnight to limit possible off-target background signaling. Cycle threshold values obtained by real-time PCR were normalized to the β-actin endogenous control and expressed as the mean log_2_ change ± SE relative to the overall lowest-expressing donor within a gene using the formula 2^−ΔΔCt^. Statistical significance was determined in the 95% confidence interval using one-way ANOVA.

## 4. Discussion

The neonatal period consistently shows disproportionately high morbidity and susceptibility to severe infection as a consequence of the immune system. Early life immune profiles present with several factors that contribute to a more suppressive state relative to adults. Among these are elevated expression of IL-27 genes and protein in human macrophages and in mice at steady-state compared to adults^16, 18, 27^. Here, we provided additional insight into potential mechanisms that regulate production of IL-27, which has a significant influence on host interactions with bacteria in early life^16, 17, 27, 45,46^. Through whole genome methyl-seq analysis, we observed that differential methylation was exclusive to the IL-27p28 promoter region between neonates and adults. While exploring hormonal influences on IL-27 production, we determined that GCs drive increases in expression of the EBI3 subunit of IL-27. Together, these data could suggest independent regulatory mechanisms for each IL-27 subunit. It is important to acknowledge that EBI3 also can pair with IL-12p35 to form the IL-35 heterodimer^47^. While our focus was to understand IL-27 regulation, IL-35 is also associated with suppressive activity^48^. IL-35 is produced primarily by regulatory B- and T-cells rather than APCs explored here, but we cannot exclude an impact of our findings on this cytokine^47, 49, 50^.

There are important implications to GC regulation of EBI3 discussed further below in the context of antenatal steroid treatment, however, this body of work demonstrated that neither methylation status of IL-27 gene promoters nor GC signals likely account for increased levels of IL-27 gene expression during steady-state conditions early in life. Specifically for GC, this suggests that endogenous levels of GC at steady-state are not consistently at adequate levels required to drive greater levels of IL-27 in murine cells of myeloid origin. Looking to alternative explanations, IL27p28 has two promoters, so it is possible that age-varied differences in cytokine expression are attributed to differential usage of the promoters during infancy versus adulthood. Assessment of publicly available sequencing data was inconclusive in addressing this possibility; however, additional studies could be conducted by exploring this hypothesis directly through ATAC-Seq, RNA-seq or the use of an inhibiting miRNA to silence each promoter and observe differences in expression.

We previously reported that progesterone treatment of human adult macrophages in vitro increased expression of IL-27p28 and EBI3, the latter more significantly^16^. Our previous report was notably different from the present body of work by using adult macrophages instead of neonatal and the use of supraphysiologic levels of progesterone, which has been shown to allow progesterone signaling through GR^51^. CBMCs, monocytes, and macrophages express GR alone (Figure 5.3) and biologically relevant concentrations of progesterone did not change IL-27 gene expression (Figure 5.4A-B). As such, we predict that in prior work progesterone was signaling promiscuously through GR instead. In this report, specificity was attributed to GR rather than PGR or a combination through pretreatment with RU486 (Figure 5.4C-D). Our findings also clearly demonstrate recruitment of GR to the EBI3 regulatory region, establishing a direct regulation of EBI3 by GC signaling. However, we cannot exclude the potential role of some indirect regulation through the control of additional transcriptional enhancers or repressors. We also cannot exclude from the prior report with adult macrophages that a membrane PR (mPR) influenced gene expression, as there is not a requirement for PR to be co-expressed or expressed to the same magnitude for mPRs to be present^52, 53^. Additional studies to explore mechanisms of progesterone signaling in neonatal macrophages could indicate another path for progesterone to influence IL-27 expression as shown previously.

Dexamethasone is commonly administered in clinical scenarios where pregnant mothers are at risk for preterm delivery. Further, corticosteroids are given to newborns experiencing respiratory distress to aid in lung development and easing the transition from ventilation devices^54^. The use of corticosteroids in early life is a debated topic due to lasting effects that follow this treatment. The weeks following birth are crucial for both the endocrine and immune system adapting to extrauterine life. The data shown here provides new insight into how treatment with corticosteroids can influence prenatal and postnatal immunity in the newborn. Previously, we have shown that IL-27 opposes host protection during neonatal sepsis^17, 20, 55^. In the present work, we show that dexamethasone can increase EBI3 expression levels. This, in turn, may lead to the unintended consequence of further compromising immunity in the newborn when presented with an infectious challenge. Understanding inherent risks that come with preterm gestation and that IL-27 has been implicated as a biomarker of sepsis severity and neonatal infection, this new data introduces new considerations for preterm interventions^56, 57^. Additional studies with preclinical models will be important to address the impact of exogenously supplied GCs and the effect on susceptibility to infection.

In summary, it is imperative to investigate the mechanisms that operate to control the unique aspects of the early life immune responses and may also promote susceptibility to infection. This will inform future intervention strategies. IL-27 remains an intriguing therapeutic target and the findings reported here have expanded our understanding of IL-27 biology early in life. Most notably, is the demonstration that GCs increase expression of the EBI3 subunit of IL-27 in macrophages, a primary cellular producer. Protocols surrounding corticosteroid usage may now also consider the impacts of this on neonatal immunity and additional safety measures to prevent severe infection.

## Data Availability Statement

All raw sequenced data are available at the Gene Expression Omnibus under accession number GSE322963. Secure token for data access during manuscript peer review can be found at opafoqoebpwrpct.

## Acknowledgments

Additional experience and support was provided by Dr. Madhavi Annamanedi, PhD, Sunilkanth Bonigala, MS, and Robert Harsh, MS. We thank Dr. Neil Billington, PhD for support in imaging protocol and analysis optimization.

## Resources and Funding

This work was supported by the NIH grants AI154129 and AI163333 awarded to CM Robinson. Additional support is provided for WVU Genomics Core Facility, WVU Bioinformatics Core (WV-INBRE P20 GM103434 and NIGMS U54 GM-104942), and WVU Microscope Imaging Facility (supported by the WVU Cancer Center and NIH P20GM121322, P20GM144230, P30GM103503, and P20GM103434).

## Authorship Contribution Statement

Conceptualization JV and CR; Design JV, JB, GH, and CR; investigation JV and CR; data curation JV, LW, JP, GH, and CR; project oversight GH and CR; original writing JV and CR; review, editing and revision JV, GH, JB, and CR.

## Consent Statement

All patients represented in this study provided prior signed informed consent.

## References

1. Sharrow D, Hug L, Liu Y, Lindt N, You D. Levels & Trends in Childhood Mortality Report 2022. UNICEF. 2022.

2. Hug L, Sharrow D, You S, Hereward M, Zhang Y. Levels and Trend in Child Mortality. 2019.

3. Basha S, Surendran N, Pichichero M. Immune responses in neonates. Expert Rev Clin Immunol. 2014;10(9):1171–84. Epub 2014/08/04. doi: 10.1586/1744666X.2014.942288. PubMed PMID: 25088080; PMCID: PMC4407563.

4. Kollmann TR, Kampmann B, Mazmanian SK, Marchant A, Levy O. Protecting the Newborn and Young Infant from Infectious Diseases: Lessons from Immune Ontogeny. Immunity. 2017;46(3):350–63. doi: 10.1016/j.immuni.2017.03.009. PubMed PMID: 28329702.

5. Caron JE, La Pine TR, Augustine NH, Martins TB, Hill HR. Multiplex Analysis of Toll-Like Receptor-Stimulated Neonatal Cytokine Response. Neonatology. 2010;97(3):266–73. doi: 10.1159/000255165. PubMed PMID: 222753613; 19955831.

6. Upham JW, Lee PT, Holt BJ, Heaton T, Prescott SL, Sharp MJ, Sly PD, Holt PG. Development of interleukin-12-producing capacity throughout childhood. Infection and immunity. 2002;70(12):6583–8. doi: 10.1128/iai.70.12.6583-6588.2002. PubMed PMID: 12438328; PMCID: PMC133015.

7. Kollmann TR, Crabtree J, Rein-Weston A, Blimkie D, Thommai F, Wang XY, Lavoie PM, Furlong J, Fortuno ES, 3rd, Hajjar AM, Hawkins NR, Self SG, Wilson CB. Neonatal innate TLR-mediated responses are distinct from those of adults. J Immunol. 2009;183(11):7150–60. Epub 2009/11/18. doi: 10.4049/jimmunol.0901481. PubMed PMID: 19917677; PMCID: PMC4556237.

8. Hornik CP, Fort P, Clark RH, Watt K, Benjamin DK, Jr., Smith PB, Manzoni P, Jacqz-Aigrain E, Kaguelidou F, Cohen-Wolkowiez M. Early and late onset sepsis in very-low-birth-weight infants from a large group of neonatal intensive care units. Early human development. 2012;88 Suppl 2(Suppl 2):S69–74. Epub 2012/05/29. doi: 10.1016/s0378-3782(12)70019-1. PubMed PMID: 22633519; PMCID: PMC3513766.

9. Weston EJ, Pondo T, Lewis MM, Martell-Cleary P, Morin C, Jewell B, Daily P, Apostol M, Petit S, Farley M, Lynfield R, Reingold A, Hansen NI, Stoll BJ, Shane AL, Zell E, Schrag SJ. The burden of invasive early-onset neonatal sepsis in the United States, 2005-2008. The Pediatric infectious disease journal. 2011;30(11):937–41. Epub 2011/06/10. doi: 10.1097/INF.0b013e318223bad2. PubMed PMID: 21654548; PMCID: PMC3193564.

10. Vink AS, Clur S-AB, Wilde AAM, Blom NA. Effect of age and gender on the QTc-interval in healthy individuals and patients with long-QT syndrome. Trends in Cardiovascular Medicine. 2018;28(1):64–75. Epub 20170803. doi: 10.1016/j.tcm.2017.07.012. PubMed PMID: 28869094.

11. Dörr HG, Heller A, Versmold HT, Sippell WG, Herrmann M, Bidlingmaier F, Knorr D. Longitudinal study of progestins, mineralocorticoids, and glucocorticoids throughout human pregnancy. The Journal of clinical endocrinology and metabolism. 1989;68(5):863–8.

12. Mastorakos G, Ilias I. Maternal and Fetal Hypothalamic-Pituitary-Adrenal Axes During Pregnancy and Postpartum. Annals of the New York Academy of Sciences. 2003;997(1):136–49. doi: 10.1196/annals.1290.016. PubMed PMID: 14644820.

13. Busada JT, Cidlowski JA. Mechanisms of Glucocorticoid Action During Development. Curr Top Dev Biol. 2017;125:147–70. Epub 20170116. doi: 10.1016/bs.ctdb.2016.12.004. PubMed PMID: 28527570.

14. Tulchinsky D, Hobel CJ, Yeager E, Marshall JR. Plasma estrone, estradiol, estriol, progesterone, and 17-hydroxyprogesterone in human pregnancy. I. Normal pregnancy. Am J Obstet Gynecol. 1972;112(8):1095–100. doi: 10.1016/0002-9378(72)90185-8. PubMed PMID: 5025870.

15. Sippell WG, Becker H, Versmold HT, Bidlingmaier F, Knorr D. Longitudinal studies of plasma aldosterone, corticosterone, deoxycorticosterone, progesterone, 17-hydroxyprogesterone, cortisol, and cortisone determined simultaneously in mother and child at birth and during the early neonatal period. I. Spontaneous delivery. J Clin Endocrinol Metab. 1978;46(6):971–85. doi: 10.1210/jcem-46-6-971. PubMed PMID: 263476.

16. Kraft JD, Horzempa J, Davis C, Jung JY, Pena MM, Robinson CM. Neonatal macrophages express elevated levels of interleukin-27 that oppose immune responses. Immunology. 2013;139(4):484–93. Epub 2013/03/08. doi: 10.1111/imm.12095. PubMed PMID: 23464355; PMCID: PMC3719065.

17. Seman BG, Vance JK, Rawson TW, Witt MR, Huckaby AB, Povroznik JM, Bradford SD, Barbier M, Robinson CM. Elevated Levels of Interleukin-27 in Early Life Compromise Protective Immunity in a Mouse Model of Gram-Negative Neonatal Sepsis. Infection and Immunity. 2020;88(3). Epub 20200220. doi: 10.1128/iai.00828-19. PubMed PMID: 31818960; PMCID: PMC7035946.

18. Jung J-Y, Gleave Parson M, Kraft JD, Lyda L, Kobe B, Davis C, Robinson J, Peña MMO, Robinson CM. Elevated interleukin-27 levels in human neonatal macrophages regulate indoleamine dioxygenase in a STAT-1 and STAT-3-dependent manner. Immunology. 2016;149(1):35–47. doi: 10.1111/imm.12625. PubMed PMID: 27238498; PMCID: PMC4981608.

19. Divens AM, Ma L, Vance JK, Povroznik JM, Hu G, Robinson CM. IL-27 producers in a neonatal BCG vaccination model are a heterogenous population of myeloid cells that are diverse in phenotype and function. Immunohorizons. 2025;9(4). doi: 10.1093/immhor/vlaf003. PubMed PMID: 40048708; PMCID: PMC11884806.

20. Vance JK, Lailler N, Divens AM, Povroznik JM, Annamanedi M, Brundage KM, Robinson CM. Interleukin-27-producing cells in gram-negative neonatal sepsis display diverse phenotypes and functions in the liver. Immunohorizons. 2025;9(8). doi: 10.1093/immhor/vlaf026. PubMed PMID: 40682363; PMCID: PMC12274645.

21. Illingworth RS, Bird AP. CpG islands--’a rough guide’. FEBS Lett. 2009;583(11):1713–20. Epub 20090418. doi: 10.1016/j.febslet.2009.04.012. PubMed PMID: 19376112.

22. Weber M, Hellmann I, Stadler MB, Ramos L, Pääbo S, Rebhan M, Schübeler D. Distribution, silencing potential and evolutionary impact of promoter DNA methylation in the human genome. Nat Genet. 2007;39(4):457–66. Epub 20070304. doi: 10.1038/ng1990. PubMed PMID: 17334365.

23. Levin ER. Rapid signaling by steroid receptors. Am J Physiol Regul Integr Comp Physiol. 2008;295(5):R1425–30. Epub 20080910. doi: 10.1152/ajpregu.90605.2008. PubMed PMID: 18784332; PMCID: PMC2584866.

24. Granner DK, Wang J-C, Yamamoto KR. Regulatory Actions of Glucocorticoid Hormones: From Organisms to Mechanisms. In: Wang J-C, Harris C, editors. Glucocorticoid Signaling: From Molecules to Mice to Man. New York, NY: Springer New York; 2015. p. 3–31.

25. Oakley RH, Cidlowski JA. The biology of the glucocorticoid receptor: New signaling mechanisms in health and disease. Journal of Allergy and Clinical Immunology. 2013;132(5):1033–44. doi: 10.1016/j.jaci.2013.09.007.

26. Graham JM. Separation of monocytes from whole human blood. ScientificWorldJournal. 2002;2:1540–3. Epub 20020607. doi: 10.1100/tsw.2002.842. PubMed PMID: 12806136; PMCID: PMC6009477.

27. Gleave Parson M, Grimmett J, Vance JK, Witt MR, Seman BG, Rawson TW, Lyda L, Labuda C, Jung JY, Bradford SD, Robinson CM. Murine myeloid-derived suppressor cells are a source of elevated levels of interleukin-27 in early life and compromise control of bacterial infection. Immunol Cell Biol. 2019;97(5):445–56. Epub 2018/12/24. doi: 10.1111/imcb.12224. PubMed PMID: 30575117; PMCID: PMC6536317.

28. Wolterink-Donselaar IG, Meerding JM, Fernandes C. A method for gender determination in newborn dark pigmented mice. Lab Animal. 2009;38(1):35–8. doi: 10.1038/laban0109-35. PubMed PMID: 19112448.

29. Annamanedi M, Vance JK, Robinson CM. Neonatal Mouse Bone Marrow Isolation and Preparation of Bone Marrow-Derived Macrophages. J Vis Exp. 2024(207). Epub 20240524. doi: 10.3791/66613. PubMed PMID: 38856198.

30. Guo W, Fiziev P, Yan W, Cokus S, Sun X, Zhang MQ, Chen P-Y, Pellegrini M. BS-Seeker2: a versatile aligning pipeline for bisulfite sequencing data. BMC Genomics. 2013;14(1):774. doi: 10.1186/1471-2164-14-774.

31. Guo W, Zhu P, Pellegrini M, Zhang MQ, Wang X, Ni Z. CGmapTools improves the precision of heterozygous SNV calls and supports allele-specific methylation detection and visualization in bisulfite-sequencing data. Bioinformatics. 2018;34(3):381–7. doi: 10.1093/bioinformatics/btx595. PubMed PMID: 28968643; PMCID: PMC6454434.

32. Kuo HC, Lin PY, Chung TC, Chao CM, Lai LC, Tsai MH, Chuang EY. DBCAT: database of CpG islands and analytical tools for identifying comprehensive methylation profiles in cancer cells. J Comput Biol. 2011;18(8):1013–7. Epub 20110108. doi: 10.1089/cmb.2010.0038. PubMed PMID: 21214365.

33. Robinson JT, Thorvaldsdóttir H, Winckler W, Guttman M, Lander ES, Getz G, Mesirov JP. Integrative genomics viewer. Nature biotechnology. 2011;29(1):24–6. doi: 10.1038/nbt.1754. PubMed PMID: 21221095; PMCID: PMC3346182.

34. Schindelin J, Arganda-Carreras I, Frise E, Kaynig V, Longair M, Pietzsch T, Preibisch S, Rueden C, Saalfeld S, Schmid B, Tinevez J-Y, White DJ, Hartenstein V, Eliceiri K, Tomancak P, Cardona A. Fiji: an open-source platform for biological-image analysis. Nature Methods. 2012;9(7):676–82. Epub 20120628. doi: 10.1038/nmeth.2019. PubMed PMID: 22743772; PMCID: PMC3855844.

35. Stirling DR, Swain-Bowden MJ, Lucas AM, Carpenter AE, Cimini BA, Goodman A. CellProfiler 4: improvements in speed, utility and usability. BMC Bioinformatics. 2021;22(1):433. Epub 20210910. doi: 10.1186/s12859-021-04344-9. PubMed PMID: 34507520; PMCID: PMC8431850.

36. Perez G, Barber GP, Benet-Pages A, Casper J, Clawson H, Diekhans M, Fischer C, Gonzalez JN, Hinrichs AS, Lee CM, Nassar LR, Raney BJ, Speir ML, van Baren MJ, Vaske CJ, Haussler D, Kent WJ, Haeussler M. The UCSC Genome Browser database: 2025 update. Nucleic Acids Res. 2025;53(D1):D1243–d9. doi: 10.1093/nar/gkae974. PubMed PMID: 39460617; PMCID: PMC11701590.

37. Wang C, Nanni L, Novakovic B, Megchelenbrink W, Kuznetsova T, Stunnenberg HG, Ceri S, Logie C. Extensive epigenomic integration of the glucocorticoid response in primary human monocytes and in vitro derived macrophages. Scientific Reports. 2019;9(1):2772. Epub 20190226. doi: 10.1038/s41598-019-39395-9. PubMed PMID: 30809020; PMCID: PMC6391480.

38. Jones MJ, Goodman SJ, Kobor MS. DNA methylation and healthy human aging. Aging Cell. 2015;14(6):924–32. Epub 20150425. doi: 10.1111/acel.12349. PubMed PMID: 25913071; PMCID: PMC4693469.

39. Martino DJ, Tulic MK, Gordon L, Hodder M, Richman TR, Metcalfe J, Prescott SL, Saffery R. Evidence for age-related and individual-specific changes in DNA methylation profile of mononuclear cells during early immune development in humans. Epigenetics. 2011;6(9):1085–94. Epub 20110901. doi: 10.4161/epi.6.9.16401. PubMed PMID: 21814035.

40. Boroditsky RS, Reyes FI, Winter JS, Faiman C. Maternal serum estrogen and progesterone concentrations preceding normal labor. Obstet Gynecol. 1978;51(6):686–91. PubMed PMID: 566408.

41. Dressing GE, Goldberg JE, Charles NJ, Schwertfeger KL, Lange CA. Membrane progesterone receptor expression in mammalian tissues: a review of regulation and physiological implications. Steroids. 2011;76(1-2):11–7. Epub 20100924. doi: 10.1016/j.steroids.2010.09.006. PubMed PMID: 20869977; PMCID: PMC3005015.

42. Cole TJ, Blendy JA, Monaghan AP, Krieglstein K, Schmid W, Aguzzi A, Fantuzzi G, Hummler E, Unsicker K, Schütz G. Targeted disruption of the glucocorticoid receptor gene blocks adrenergic chromaffin cell development and severely retards lung maturation. Genes Dev. 1995;9(13):1608–21. doi: 10.1101/gad.9.13.1608. PubMed PMID: 7628695.

43. Lockwood CJ, Radunovic N, Nastic D, Petkovic S, Aigner S, Berkowitz GS. Corticotropin-releasing hormone and related pituitary-adrenal axis hormones in fetal and maternal blood during the second half of pregnancy. J Perinat Med. 1996;24(3):243–51. doi: 10.1515/jpme.1996.24.3.243. PubMed PMID: 8827573.

44. Cadepond F, Ulmann A, Baulieu EE. RU486 (mifepristone): mechanisms of action and clinical uses. Annu Rev Med. 1997;48:129–56. doi: 10.1146/annurev.med.48.1.129. PubMed PMID: 9046951.

45. Povroznik JM, Wang L, Annamanedi M, Bare RL, Akhter H, Hu G, Robinson CM. The influence of interleukin-27 on metabolic fitness in a murine neonatal model of bacterial sepsis. American Journal of Physiology-Endocrinology and Metabolism. 2025;0(0):null. Epub 20250114. doi: 10.1152/ajpendo.00243.2024. PubMed PMID: 39810405; PMCID: PMC12184839.

46. Povroznik JM, Akhter H, Vance JK, Annamanedi M, Dziadowicz SA, Wang L, Divens AM, Hu G, Robinson CM. Interleukin-27-dependent transcriptome signatures during neonatal sepsis. Frontiers in Immunology. 2023;14:1124140. Epub 20230220. doi: 10.3389/fimmu.2023.1124140. PubMed PMID: 36891292; PMCID: PMC9986606.

47. Devergne O, Birkenbach M, Kieff E. Epstein-Barr virus-induced gene 3 and the p35 subunit of interleukin 12 form a novel heterodimeric hematopoietin. Proceedings of the National Academy of Sciences of the United States of America. 1997;94(22):12041–6. doi: 10.1073/pnas.94.22.12041. PubMed PMID: 9342359; PMCID: PMC23696.

48. Song M, Ma X. The Immunobiology of Interleukin-35 and Its Regulation and Gene Expression. Adv Exp Med Biol. 2016;941:213–25. doi: 10.1007/978-94-024-0921-5_10. PubMed PMID: 27734415.

49. Bardel E, Larousserie F, Charlot-Rabiega P, Coulomb-L’Herminé A, Devergne O. Human CD4+ CD25+ Foxp3+ regulatory T cells do not constitutively express IL- 35. J Immunol. 2008;181(10):6898–905. doi: 10.4049/jimmunol.181.10.6898. PubMed PMID: 18981109.

50. Shen P, Roch T, Lampropoulou V, O’Connor RA, Stervbo U, Hilgenberg E, Ries S, Dang VD, Jaimes Y, Daridon C, Li R, Jouneau L, Boudinot P, Wilantri S, Sakwa I, Miyazaki Y, Leech MD, McPherson RC, Wirtz S, Neurath M, Hoehlig K, Meinl E, Grützkau A, Grün JR, Horn K, Kühl AA, Dörner T, Bar-Or A, Kaufmann SHE, Anderton SM, Fillatreau S. IL-35-producing B cells are critical regulators of immunity during autoimmune and infectious diseases. Nature. 2014;507(7492):366–70. Epub 20140223. doi: 10.1038/nature12979. PubMed PMID: 24572363; PMCID: PMC4260166.

51. Hierweger AM, Engler JB, Friese MA, Reichardt HM, Lydon J, DeMayo F, Mittrücker HW, Arck PC. Progesterone modulates the T-cell response via glucocorticoid receptor-dependent pathways. Am J Reprod Immunol. 2019;81(2):e13084. Epub 20190128. doi: 10.1111/aji.13084. PubMed PMID: 30604567; PMCID: PMC7457140.

52. Dressing GE, Alyea R, Pang Y, Thomas P. Membrane progesterone receptors (mPRs) mediate progestin induced antimorbidity in breast cancer cells and are expressed in human breast tumors. Horm Cancer. 2012;3(3):101–12. doi: 10.1007/s12672-012-0106-x. PubMed PMID: 22350867; PMCID: PMC10358000.

53. Lacroix M, Leclercq G. Relevance of breast cancer cell lines as models for breast tumours: an update. Breast Cancer Res Treat. 2004;83(3):249–89. doi: 10.1023/B:BREA.0000014042.54925.cc. PubMed PMID: 14758095.

54. Boscarino G, Cardilli V, Conti MG, Liguori F, Repole P, Parisi P, Terrin G. Outcomes of postnatal systemic corticosteroids administration in ventilated preterm newborns: a systematic review of randomized controlled trials. Front Pediatr. 2024;12:1344337. Epub 20240214. doi: 10.3389/fped.2024.1344337. PubMed PMID: 38419972; PMCID: PMC10899705.

55. Povroznik JM, Robinson CM. IL-27 regulation of innate immunity and control of microbial growth. Future Sci OA. 2020;6(7):FSO588-FSO. Epub 20200617. doi: 10.2144/fsoa-2020-0032. PubMed PMID: 32802395; PMCID: PMC7421895.

56. He Y, Du Wx, Jiang Hy, Ai Q, Feng J, Liu Z, Yu Jl. Multiplex Cytokine Profiling Identifies Interleukin-27 as a Novel Biomarker For Neonatal Early Onset Sepsis. Shock. 2017;47(2):140–7. doi: 10.1097/shk.0000000000000753. PubMed PMID: 27648693.

57. Wong HR, Cvijanovich NZ, Hall M, Allen GL, Thomas NJ, Freishtat RJ, Anas N, Meyer K, Checchia PA, Lin R, Bigham MT, Sen A, Nowak J, Quasney M, Henricksen JW, Chopra A, Banschbach S, Beckman E, Harmon K, Lahni P, Shanley TP. Interleukin-27 is a novel candidate diagnostic biomarker for bacterial infection in critically ill children. Critical care (London, England). 2012;16(5):R213. Epub 2012/10/31. doi: 10.1186/cc11847. PubMed PMID: 23107287; PMCID: PMC3682317.

