## Supplemental Material for "Glucocorticoid signaling regulates expression of the EBI3 subunit of IL-27 in neonatal macrophages: Implications for antenatal corticosteroid therapy"

**Supplemental Materials**

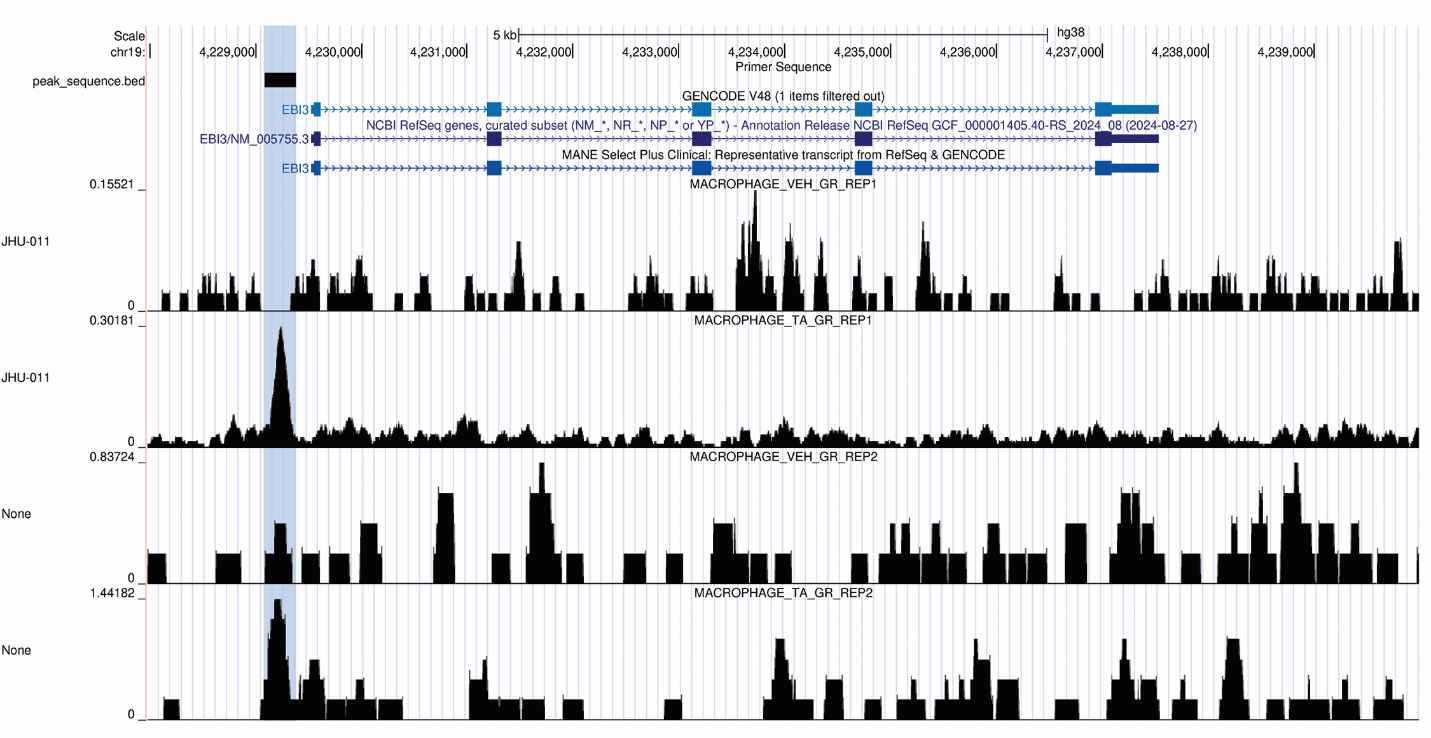

*Figure 6.1 Primer design in UCSC genome browser.*

Publicly available ChIP-Seq data from Wang, 2019^37^ was visualized using the UCSC Genome Browser to identify GR binding in the EBI3 promoter region in the JHU-011 cell line (top two tracks) and PBMCs (bottom two tracks) from public datasets. Shown on the Y-axis are normalized ChIP-Seq read density. Shown is each cell type treated with and without triamcinolone, a synthetic GC. Primers were designed for the highlighted blue region containing the GR binding peak.

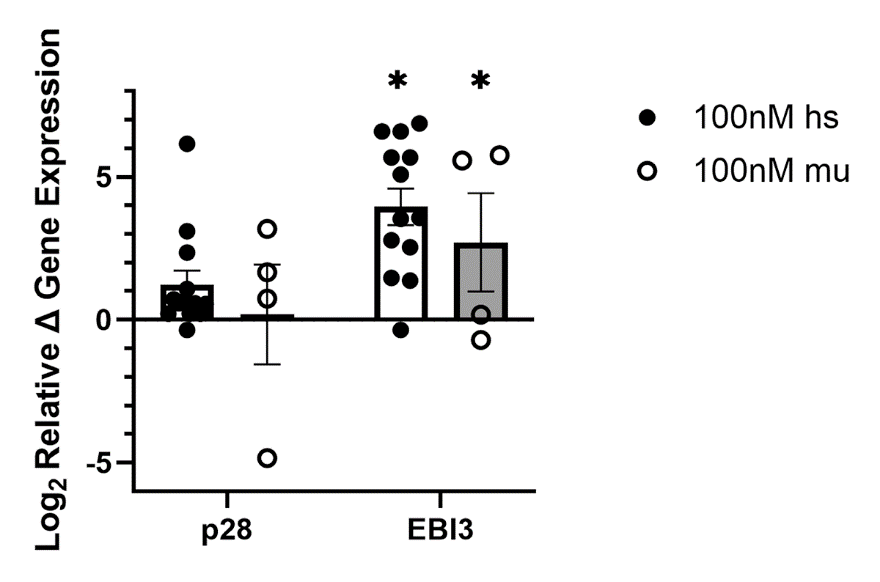

*Figure 6.2 Human and murine neonatal macrophages present similar gene expression profiles in response to dexamethasone stimulation.*

Neonatal macrophages were differentiated from either CBMCs (hs) or bone marrow (mu) and stimulated with dexamethasone at 100 nM for 48 h. Cells were placed in 1% serum containing media overnight to limit possible off-target background signaling. Values were normalized to the GAPDH (human) or β-actin (murine) endogenous control and expressed as the log_2_ change relative to vehicle controls using 2^−∆∆Ct^. Mean log_2_ fold change ± SEM was analyzed for statistical significance in the 95% confidence interval by one-way ANOVA.
